# Whole-Genome Sequence Typing shows extensive diversity of *Listeria monocytogenes* in the outdoor environment and poultry processing plants

**DOI:** 10.1101/2020.06.18.160705

**Authors:** Swarnali Louha, Richard J. Meinersmann, Zaid Abdo, Mark E. Berrang, Travis C. Glenn

## Abstract

A reliable and standardized classification of *Listeria monocytogenes* (*Lm*) is important for accurate strain identification during outbreak investigations. Current whole-genome sequencing (WGS) based approaches for strain characterization either lack standardization, rendering them less suitable for data exchange, or are not freely available. Thus, we developed a portable and open-source tool Haplo-ST to improve standardization and provide maximum discriminatory potential to WGS data tied to an MLST (multi locus sequence typing) framework. Haplo-ST performs whole-genome MLST (wgMLST) for *Lm* while allowing for data exchangeability worldwide. This tool takes in (i) raw WGS reads as input, (ii) cleans the raw data according to user specified parameters, (iii) assembles genes across loci by mapping to genes from reference strains, (iv) assigns allelic profiles to assembled genes and provides a wgMLST subtyping for each isolate. Data exchangeability relies on the tool assigning allelic profiles based on a centralized nomenclature defined by the widely-used BIGSdb-*Lm* database. Tests on Haplo-ST’s performance with simulated reads from *Lm* reference strains yielded a high sensitivity of 97.5%, and coverage depths of ≥ 20× was found to be sufficient for wgMLST profiling. We used Haplo-ST to characterize and differentiate between two groups of *Lm* isolates, derived from the natural environment and poultry processing plants. Phylogenetic reconstruction showed sharp delineation of lineages within each group and no lineage-specificity was observed with isolate phenotypes (transient vs. persistent) or origins. Genetic differentiation analyses between isolate groups identified 21 significantly differentiated loci, potentially enriched for adaptation and persistence of *Lm* within poultry processing plants.

**IMPORTANCE:** We have developed an open-source tool that provides allele-based subtyping of *Lm* isolates at the whole genome level. Along with allelic profiles, this tool also generates allele sequences, and identifies paralogs, which is useful for phylogenetic tree reconstruction and deciphering relationships between closely related isolates. More broadly, Haplo-ST is flexible and can be adapted to characterize the genome of any haploid organism simply by installing an organism-specific gene database. Haplo-ST also allows for scalable subtyping of isolates; fewer reference genes can be used for low resolution typing, whereas higher resolution can be achieved by increasing the number of genes used in the analysis. Our tool enabled clustering of *Lm* isolates into lineages and detection of potential loci for adaptation and persistence in food processing environments. Findings from these analyses highlights the effectiveness of Haplo-ST in subtyping and evaluating relationships among isolates for routine surveillance, outbreak investigations and source tracking.

## INTRODUCTION

*Listeria monocytogenes* is an opportunistic foodborne pathogen associated with significant public health concern worldwide, with an estimated 1600 illnesses and 260 deaths occurring annually (1, 2) and an estimated annual economic burden of $2.8 billion in the United States (3). *Lm* primarily causes the food borne illness listeriosis but may also cause septicemia, encephalitis and meningitis in the immunocompromised, newborn and elderly and severe complications in pregnancies leading to stillbirths and miscarriages (4). *Lm* is ubiquitous in the natural environment. Although all environmental isolates of *Lm* have the potential to cause disease, it is uncertain if some clones are more virulent than others (5).

Listeriosis mainly occurs through the consumption of food such as meat, fish and dairy products which become contaminated in food processing facilities during manufacturing, post-processing or storage for extended periods of time before consumption (6). Nearly all sporadic and epidemic human listeriosis cases have been linked to contaminated food or feed (7). Within food processing facilities, *Lm* can adapt to survive conditions used for food preservation and safety; it can replicate at low temperatures, high salt conditions and withstand disinfectants and nitrate preservation methods. These, together with the ability to form biofilms on food contact surfaces can facilitate persistence of *Lm* in food facilities (7). In fact, studies have shown that *Lm* can persist for more than 10 years in food processing facilities (8, 9). Persistence may also arise from the survival of the bacteria in nooks not reached by regular cleaning and sanitation procedures. Often, this results in cross-contamination of the final product multiple times, which increases the risk of an outbreak. On the other hand, frequent introduction of *Lm* from external sources may result in a high prevalence of transient strains within food facilities (10). Contaminating strains of *Lm* are later released from food facilities into the natural environment via effluents (11, 12). Hence, food regulatory authorities should implement effective surveillance and control measures to discriminate between transient and persistent strains, decrease harborage and prevent dissemination of *Lm*. Additionally, it is important to investigate the relatedness of strains of *Lm* involved in a single contamination event for accurate source tracking. Such investigations can help optimize effective control measures to prevent recurrence of contamination in food processing facilities (10).

Molecular subtyping techniques have been traditionally used for strain discrimination and identification of degrees of genetic relatedness among isolates (13). While many other subtyping methods (ribotyping, REP-PCR, MLEE) have been used in the past, pulsed-field gel electrophoresis (PFGE) has been the gold standard subtyping tool for *Lm* for many years (14). Although PFGE has been extremely useful in outbreak investigations and source tracking of *Lm* at food settings (10), it is time consuming, labor-intensive, expensive, and difficult to standardize (15, 16). Moreover, it provides little information on the genetic variation within or phylogenetic relationships among strains, limiting our overall understanding of evolutionarily important traits such as virulence. In contrast, sequence-based approaches are promising tools for strain typing and phylogeny assessment (17). Multi-locus sequence typing (MLST) differentiates strains by detecting variation within the nucleotide sequences of seven housekeeping genes. Every isolate is defined by a sequence type (ST), which consists of a combination of seven allelic profiles. Groups of STs sharing a minimum of six identical alleles along with an ST acting as the ‘central genotype’ forms clonal complexes (CCs), which can be geographically and temporally widespread (17). Conventional MLST has been used to describe the population structure of *Lm*, and has shown that *Lm* forms a structured population consisting of four divergent lineages (I-IV) (17, 18). Each lineage is comprised of multiple serotypes; with lineage I containing serotypes 1/2b, 3b, 4b, 4d, 4e and 7; lineage II, serotypes 1/2a, 1/2c, 3a, 3c; lineage III: serotypes 1/2a, 4a, 4b and 4c; and lineage IV: 4a and 4c. About 96% of all human listeriosis cases are caused by Lineage I and II; serotypes 1/2a, 1/2b and 4b (7). Lineage I strains are known to be highly clonal, indicating strong selection of genetic traits of fitness within the host, whereas Lineage II strains show higher rates of recombination than Lineage I and this increased genome plasticity may help in adapting to diverse ecological niches (19, 20). This is supported by the fact that Lineage I strains are predominantly linked to human clinical infection and animal listeriosis, whereas Lineage II strains are more commonly associated with food contamination and the environment. Lineage III and IV strains occur less frequently among humans and have been linked to animals (21).

The advent of next generation sequencing technologies has facilitated whole genome sequencing (WGS)-based subtyping at low costs and speeds exceeding that of traditional MLST. WGS enables easy availability of total bacterial genomes that allow strain discrimination at very high resolution. WGS also provides the ability to infer phylogenetic relationships among isolates, along with access to additional information such as virulence and resistance markers (6). WGS-based subtyping has become extremely valuable for epidemiological surveillance, outbreak detection and source tracking in the United States (22), France (23), Germany (24), Denmark (25), and Australia (26) among other countries. WGS-based subtyping approaches are either based on single nucleotide polymorphisms (SNPs) (22, 27), or on gene-by-gene allelic profiling of a defined set of genes in the genome (10, 23). Although studies have shown that both SNP-based subtyping and whole-genome based allelic profiling show similar discriminatory power and clustering among isolates (10, 28), SNP-based approaches are dependent on the choice of a reference genome, can be difficult to interpret, and are limited to assessing closely related isolates (10). These limitations are overcome by gene-by-gene approaches, which are based on allelic variation of a predefined set of genes from either the core genome (cgMLST) or on a set of genes from both core and accessory genome (wgMLST).

Several cgMLST schemes have been developed for subtyping *Lm* (15, 29-31). These cgMLST schemes are different from each other with respect to the method employed, the diversity and number of isolates used in scheme development, and the number of loci used in each scheme. These differences between cgMLST schemes can impact communication on cluster detection between different laboratories, as knowledge on the type of core genome scheme, assembler, assembler version, and sequencing technology used for cluster detection becomes crucial (32). Further, cgMLST finds differences only within the core genome of *Lm*, which represents ∼58% of the genome in terms of number of genes and ∼54% in terms of the length of the genome. Though this level of differentiation may be sufficient for discriminating outbreak strains from epidemiologically unrelated strains, investigating persistence and source tracking of root-cause analysis requires increased discriminatory power beyond cgMLST (10). These problems can be addressed with a standardized wgMLST-based subtyping, which can profile allelic differences among *Lm* strains on a genome-wide scale.

In this study, we present the Haploid Sequence-Typer (Haplo-ST), a tool that can perform wgMLST for *Lm* while allowing for data exchangeability worldwide. In contrast to the commercial genome-wide MLST scheme developed by BioNumerics^®^ (Applied Maths NV, Belgium) and being used by the US CDC and PulseNet International, Haplo-ST is open-source. Haplo-ST takes in WGS reads as input, assembles genes across loci by mapping to genes from reference strains and assigns allelic profiles to the assembled genes, thus providing a wgMLST profile for each isolate sequenced. The use of an allelic nomenclature defined by the widely-used Institute Pasteur BIGSdb-*Lm* database (available at http://bigsdb.pasteur.fr/listeria) facilitates standardized genotyping and easy inter-laboratory data exchange. Further, we conducted *in silico* tests on the sensitivity of Haplo-ST and evaluated accuracy of whole-genome sequence typing with varying levels of sequence coverage.

After developing Haplo-ST, we used it to characterize and differentiate between two groups of *Lm* isolates; the first group was obtained from the natural environment and the second group was obtained from poultry processing plants. Isolates from the natural environment were sampled from agricultural sites, forests, sites impacted by water pollution control plants (WPCP) and mixed-use sites. Because Lineage III strains are mostly associated with animals (21), we hypothesized that the majority of isolates obtained from agricultural/pastoral sites would belong to lineage III. Secondly, isolates obtained from the poultry processing plants contained both transient and persistent strains of *Lm*. Because strains belonging to lineage II are predominantly associated with contaminated food, we hypothesized that most of the strains isolated from the poultry processing plants would belong to lineage II. Previous research has shown that persistent strains have increased adhesion and biofilm formation capacity (33) and are genetically distinct from transient strains (34). However, larger-scale studies of the extent of genetic variation existing between persistent and transient strains are still needed. Furthermore, understanding the genetic diversity between *Lm* isolates present in the natural environment and food processing plants can indicate specific traits selected in the processing plant environment, and the genetic and physiological factors responsible for the persistent phenotype.

This study aims to (i) develop Haplo-ST for performing wgMLST of *Lm* isolates; (ii) establish phylogenetic relationships within the two group of *Lm* isolates obtained from the outdoor environment and poultry processing plants; (iii) examine if there exists any lineage-specific association of isolates obtained from (a) different sites in the natural environment, and (b) transient and persistent strains; (iv) analyze the extent of genetic variation between (a) isolates obtained from the natural environment and poultry processing plants, and (b) transient and persistent strains of *Lm*. We describe below how we achieved these aims.

## RESULTS

### Sensitivity of Haplo-ST

Allelic profiles derived from Haplo-ST for *Lm* strains EGD-e and 4b F2365 were compared to allele profiles of 1826 loci in EGD-e and 1825 loci in 4b F2365 respectively. On average, 4.4% of genes had uncalled alleles; this may be due to the inability of short reads to assemble these genes completely. Amongst the loci that were assigned allele designations, reproducibility of allele calls with Haplo-ST was significant, yielding an average sensitivity of 97.5% over eight simulated datasets for coverage depths of ∼ 80× (Phred quality score ≥ 20 for ≥ 90% bases in the retained reads).

### Dependency of Haplo-ST on sequencing depth

The number of genes correctly profiled by Haplo-ST increased rapidly from a sequencing depth of 5× to 10×, then increased modestly from 10× to 20× and did not increase further beyond a depth of 20× (Fig. 2A). The number of genes assigned an erroneous allele ID (i.e., misassigned) and the number of genes missing an allele ID assignment (i.e., missing or uncalled alleles) decreased significantly up to a depth of 20×, improved slightly at 30× and then remained stable at higher sequencing depths (Fig. 2B). The average number of genes partially assembled by YASRA and thus giving rise to uncalled alleles by BIGSdb-*Lm* remained similar over all sequencing depths. This may be due to the presence of low complexity regions within these genes which could not be sequenced with short Illumina reads. From these results we conclude that sequencing depths ≥ 20× will perform well in Haplo-ST for wgMLST profiling of *Lm* isolates.

**Figure 1:**
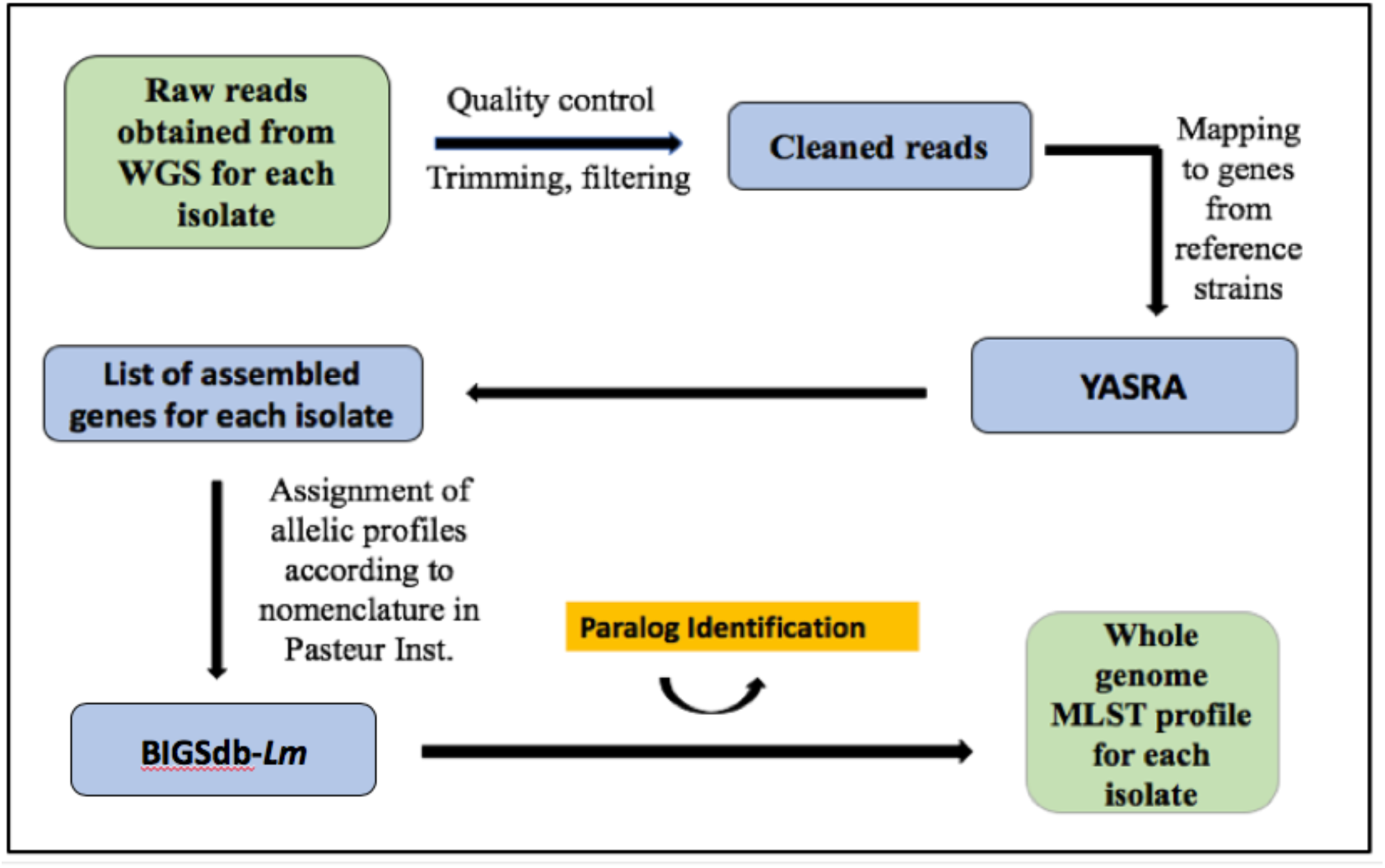
Haplo-ST, a tool for wgMLST profiling of *Listeria monocytogenes* (*Lm*) from WGS reads.

**Figure 2:**
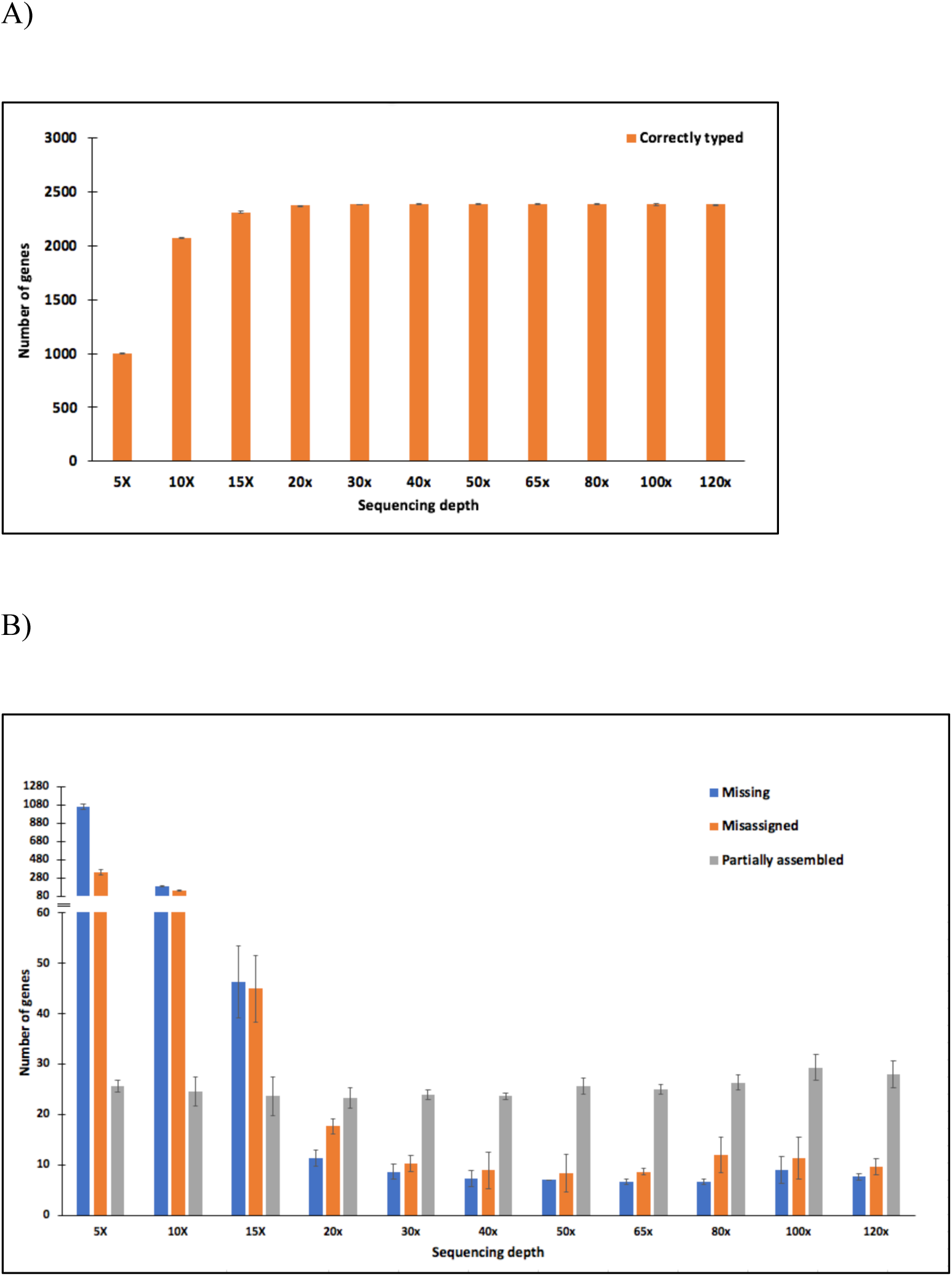
Dependency of Haplo-ST on sequencing depth. (a) Simulation of the number of genes correctly profiled by Haplo-ST across sequencing depths ranging from 5× - 120× (b) The number of genes missing an allele ID assignment, the number of genes misassigned an erroneous allele ID and the number of genes partially assembled with Haplo-ST across different sequencing depths.

### wgMLST profiling of Lm isolates

Haplo-ST generated a wgMLST profile of each *Lm* isolate from WGS reads (File S7). A list of assembled gene sequences identified in each isolate were also provided by Haplo-ST (available at https://bit.ly/3e9KM6g).

### Identification of paralogs

We used two different approaches to identify paralogous genes in our dataset. With our first approach, Haplo-ST generated a list of paralogous genes for each *Lm* isolate while profiling isolates. Our second approach identified 133 paralogous genes (File S3) in BIGSdb-*Lm.* Comparison of the two approaches for paralog detection showed that BIGSdb-*Lm* correctly identifies all paralogous genes. However, in a few instances BIGSdb-*Lm* incorrectly identifies non-paralogous genes as ‘exact matches’ to each other (File S4). On further examination, we found that in such cases, two allele sequences partially matched across their lengths with a 100% identity (see examples in File S4).

### Phylogenetic relationships among *Lm* isolates

*Lm* isolates obtained from the natural environment formed three distinct clusters in the phylogenetic tree (Fig. 3A), with each cluster containing a specific lineage (I, II and III) of *Lm* strains. A majority of the isolates belonged to lineage II (68%), followed by isolates from lineage III (17%), lineage I (5%), and 15 isolates (9%) that could not be genotyped into lineages with lineage-specific probes clearly clustered in lineage II. A few isolates were distantly related to these clusters and an assembly of the 16s rRNA sequence of these isolates showed that they belonged to non-pathogenic *Listeria* species, *L. seeligeri* (n=2) and *L. welshimeri* (n=1). Isolates from the three *Lm* lineages were found to be randomly distributed across the four sampling locations (Fig. 3B). This was confirmed with Fisher’s exact test (*α* = 0.05) which failed to show any lineage-specific association of isolates with sampling sites (*P* = 0.067). Isolates sampled from pastoral sites contained a mix of lineages I, II and III with a majority of isolates belonging to lineage II (86%) thus failing to support our hypothesis that isolates sampled from the pastoral sites would mostly belong to lineage III.

**Figure 3:**
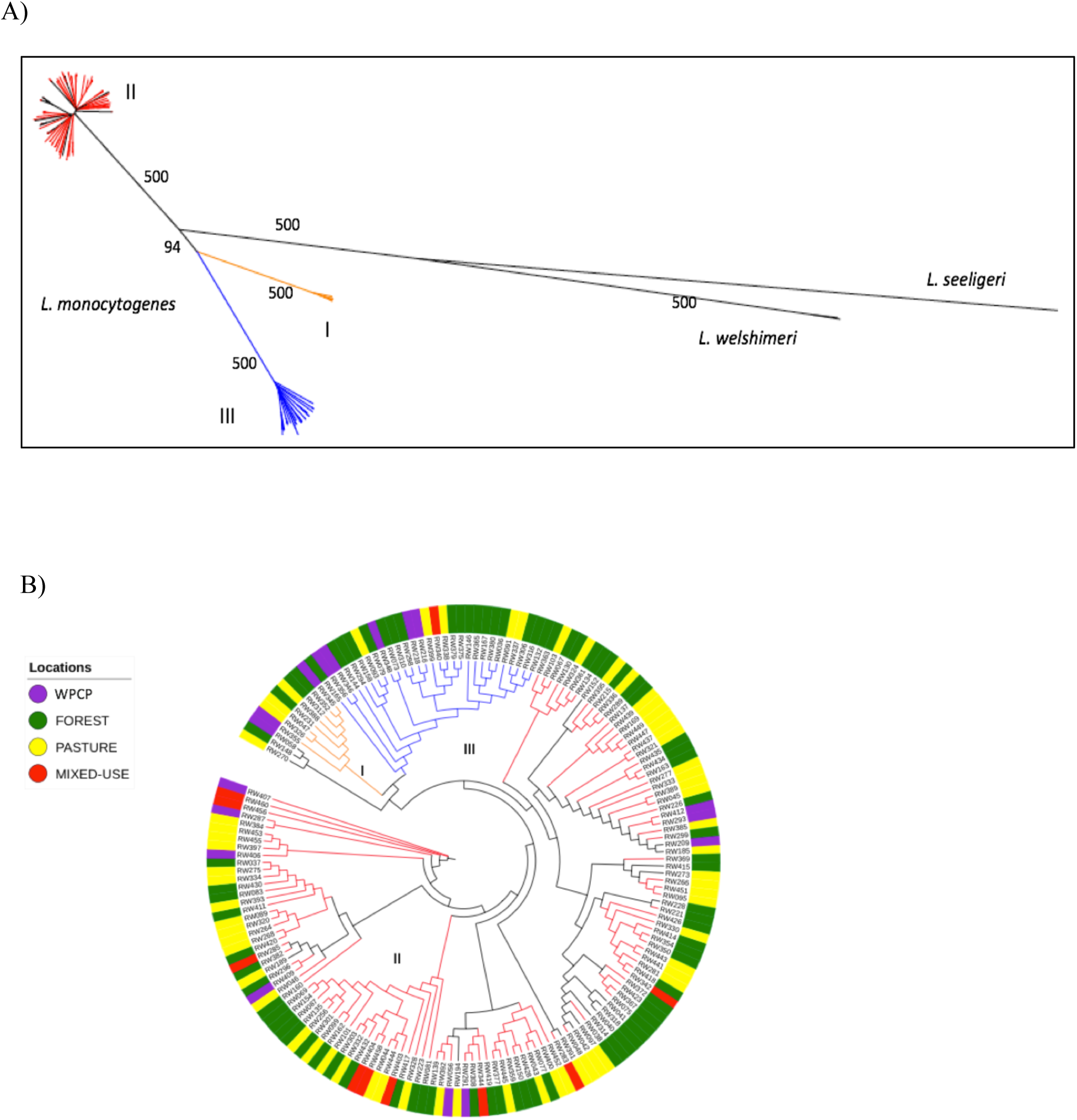
Phylogenetic relationships between isolates collected from the natural environment. (3a) *Lm* isolates belonging to lineages I (orange), II (red) and III (blue) form separate clusters in the phylogenetic tree. (3b) Random distribution of three lineages of *Lm* found at different sampling sites.

The phylogenetic tree constructed from isolates obtained from poultry processing plants had two major clusters (Fig. 4A); one containing isolates belonging to lineage I (35%) and the other containing lineage II (59%) isolates. Twelve isolates could not be classified into lineages by genotyping with lineage-specific probes; of these 12, 3 isolates clustered in lineage I and 4 isolates clustered in lineage II. The remaining 5 isolates were distantly related from the two major lineages in the tree and were identified as non-pathogenic species of *Listeria, L. innocua* (n=4) and *L. welshimeri* (n=1). Persistent strains were more abundant (65%) than transient strains (35%) and correlation of *Lm* lineages with transient vs. persistent phenotypes was not significant (Fishers exact test, *P* = 0.86) (Fig. 4B).

**Figure 4:**
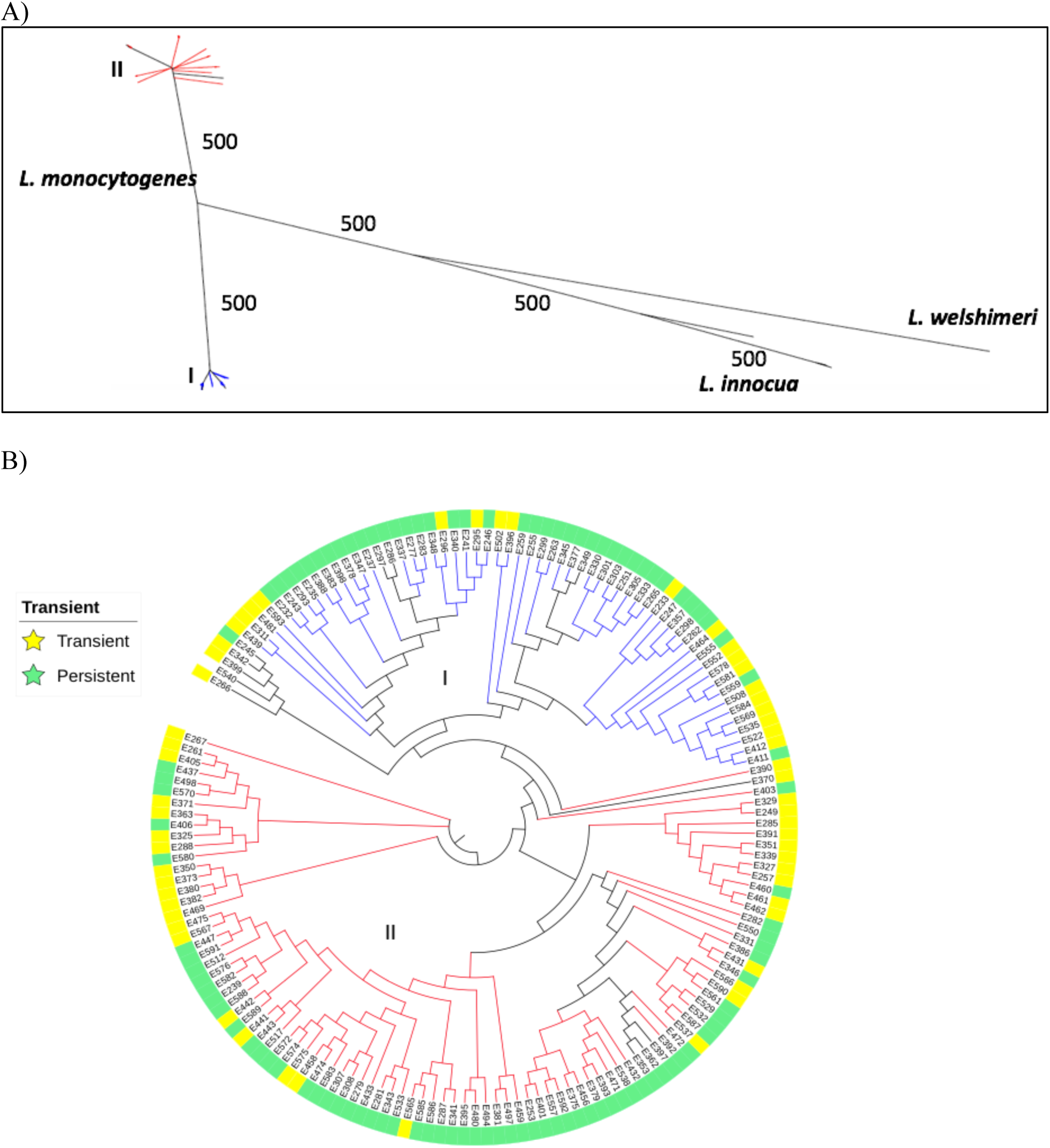
Phylogenetic relationships between isolates collected from poultry processing plants. (4a) *Lm* lineages I (blue) and II (red) form two separate groups in the phylogenetic tree, with the majority of isolates belonging to lineage II. (4b) Persistent strains were more abundant than transient strains, but there was no lineage-specific association of persistent/transient strains.

### Analysis of molecular variance

The wgMLST profiles were filtered for paralogous genes and assigned custom allele ID’s for new alleles (File S8). The results from AMOVA showed that most genetic variation was contained within isolates obtained from the natural environment and poultry processing plants (91%), with only 9% attributed to variation between the two groups (Table 1). To detect loci with significant genetic variation between the two groups, we calculated population specific F_ST_ values for each locus separately with locus-by-locus AMOVA. We chose 111 loci (top 5% of F_ST_ distribution; F_ST_ ≥ 0.149) with the highest F_ST_ values as loci having considerable genetic variation between isolates obtained from the natural environment and poultry processing plants (Fig. 5A, File S5). Additionally, results from AMOVA considering only isolates from the poultry processing plants suggested that majority of the genetic variance was within isolates (96.18%), and the remaining variation (3.18%) was between the transient and persistent groups of strains (Table 1). In this case, 102 loci (upper 5%; F_ST_ ≥ 0.782) were identified as having the most divergence between the transient and persistent strains (Fig. 5B, File S6). A set of 21 loci were common among the loci with highest F_ST_ values in both levels of AMOVA (ie. 111 loci in natural environment vs. poultry processing plants, and 102 loci in transient vs. persistent), and might play a role in the adaptation and persistence of *Lm* in poultry processing environments (Table 2).

**Table 1:** Global AMOVA results weighted over all variable loci in the two groups of *L. monocytogenes* isolates.

| Groups of isolates | Source of variation | Variance components | Variation (%) | Fixation index |
| --- | --- | --- | --- | --- |
| Natural Environment<br>vs. Poultry Processing<br>plants | Among groups | 86.61 | 9.00 | $F_{ST} =$<br>0.09002 |
|  | Within groups | 875.51 | 91.00 |  |
| Persistent vs.<br>Transient | Among groups | 32.64 | 3.82 | $F_{ST} =$<br>0.03821 |
|  | Within groups | 821.66 | 96.18 |  |

**Table 2:** Genes showing significant genetic differentiation between groups of *Lm* isolates collected from the natural environment vs. poultry processing plants and transient vs. persistent strains and may be enriched for adaptation and persistence of *Lm* in poultry processing environments.

| Gene name | Gene product (obtained from RefSeq) | Biological Function (obtained from KEGG) |
| --- | --- | --- |
| lmo0875 | PTS beta-glucoside transporter subunit IIB | Carbohydrate metabolism, Membrane transport |
| lmo1949 | hypothetical protein | Ribosome biogenesis |
| lmo2650 | MFS transporter | Carbohydrate metabolism, Membrane transport |
| lmo1210 | hypothetical protein | Electrochemical potential-driven transporters |
| lmo0687 | hypothetical protein | Peptidase |
| lmo0694 | hypothetical protein | Unknown function |
| lmo0964 | hypothetical protein (thioredoxin) | Oxidative stress, Signaling |
| lmo2078 | hypothetical protein | Transfer RNA biogenesis |
| lmo2383 | monovalent cation/H<sup>+</sup> antiporter subunit F | Electrochemical potential-driven transporters |
| lmo2548 | 50S ribosomal protein L31 | Translation |
| lmo1776 | hypothetical protein | Unknown function |
| lmo1960 | ferrichrome ABC transporter<br>ATP-binding protein | Iron complex transporter |
| lmo1336 | 5-formyltetrahydrofolate cyclo-<br>ligase | Metabolism of cofactors and vitamins |
| lmo2689a | hypothetical protein | Uncharacterized |
| lmo1294 | tRNA delta(2)-<br>isopentenylpyrophosphate<br>transferase | Transfer RNA biogenesis, Biosynthesis of<br>secondary metabolites |
| lmo2640 | hypothetical protein | Metabolism of terpenoids and polyketides,<br>Biosynthesis of secondary metabolites |
| lmo0360 | DeoR family transcriptional<br>regulator | Unknown function |
| lmo1817 | hypothetical protein | Metabolism of cofactors and vitamins |
| lmo2073 | ABC transporter ATP-binding<br>protein | Translation factor |
| lmo1205 | cobalamin biosynthesis protein<br>CbiN | Membrane transport |
| lmo1464 | diacylglycerol kinase | Glycan biosynthesis and metabolism |

**Fig 5:**
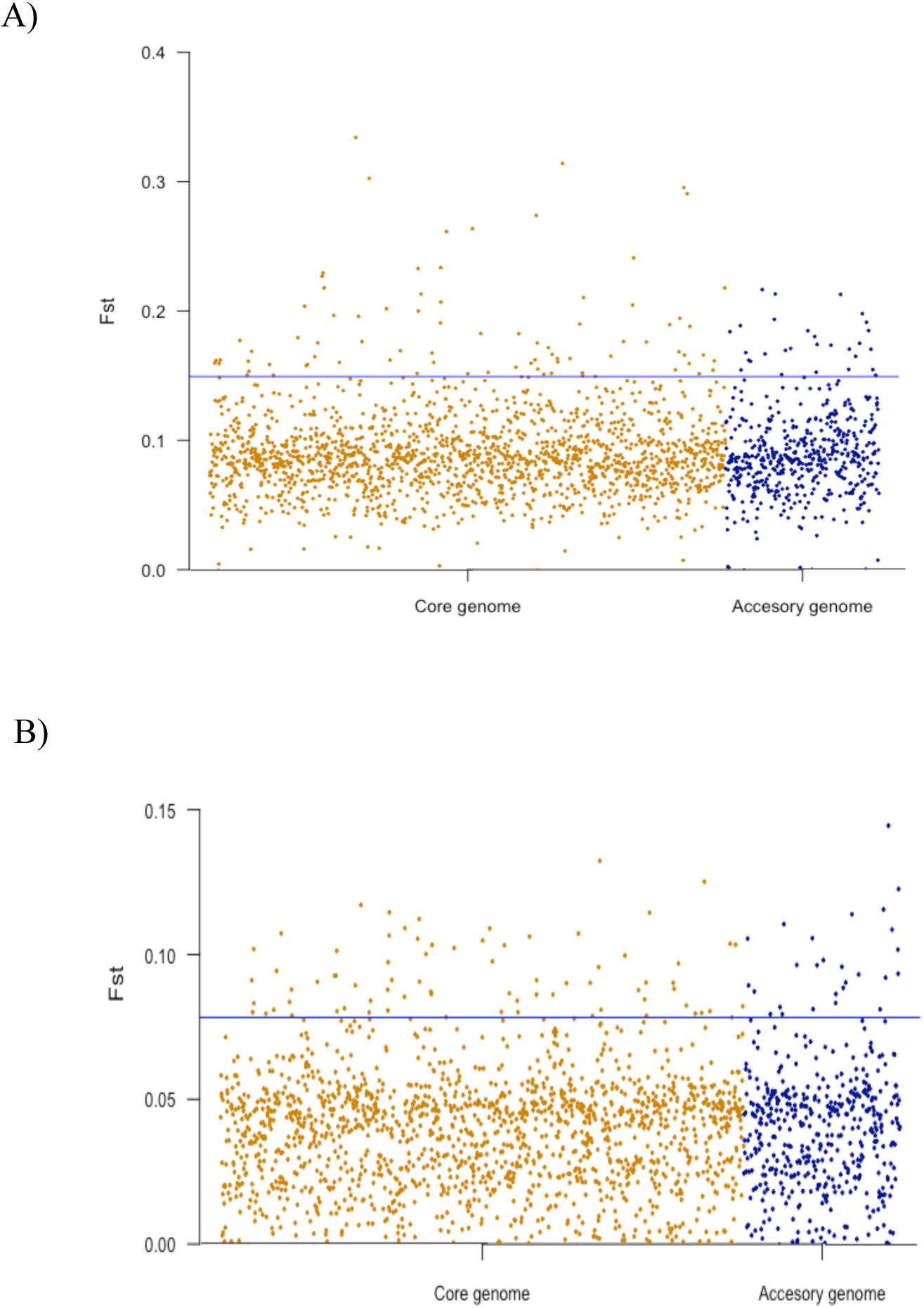
Manhattan plots of genome-wide F_ST_ values between (a) *Lm* isolates obtained from the natural environment and poultry processing plants (b) groups of transient and persistent *Lm* strains. F_ST_ values are shown on the y axis. The loci are arranged in two groups on the x axis; the first group consisting of loci present in the core genome and the other group consisting of loci in the accessory genome as specified in BIGSdb-*Lm.* The significant thresholds (blue line) are set at the top 5% of the F_ST_ distribution.

## DISCUSSION

Molecular characterization of *Listeria monocytogenes* is important for outbreak detection, surveillance, epidemiological studies and in the development of effective control strategies for listeriosis. We have developed a freely available and portable tool, Haplo-ST, that can be used for wgMLST profiling of *Lm* from WGS data. Our tool uses the centralized nomenclature of *Lm* genotypes publicly accessible in the BIGSdb-*Lm* database and the BIGSdb software for calling alleles, which facilitates sharing and comparing data between public health laboratories worldwide, and enables international source tracking and outbreak investigation. We have shown that the reproducibility of allele calls by Haplo-ST has high sensitivity (error rate ∼ 2.5%), and sequencing depths of ∼20× are sufficient for assembling alleles. Because our genotyping technique assembles alleles directly from WGS data by mapping to corresponding reference genes before allele typing, it is computationally faster and less error-prone than other subtyping techniques that require *de-novo* assembly of genomes prior to allele identification and subtyping (15, 31). This property also allows for the scalable characterization of isolates based on the needs of the researcher, as some questions require more discrimination among isolates than others. For example, lower resolution is required for assignment of isolates to a specific lineage or clonal complex whereas higher levels of discrimination are needed for outbreak detection and investigation of within-patient variations (35). In this regard, Haplo-ST is flexible because it can be used with custom sets of fewer reference genes for low resolution typing, whereas higher resolution can be achieved by increasing the number of reference genes used in the analysis. The time required for low resolution typing is low and increases with the increase in typing resolution. For example, on a system with a quad-core processor running at 3.6 GHz and 50 GB of RAM, the time taken for subtyping 100, 500 and 1000 loci was 1.4, 6.2, and 12.8 hrs respectively.

The motivation to develop Haplo-ST was to design an open-source platform which can automate the translation of Illumina WGS data from *Lm* collections into their corresponding wgMLST genotypes, thereby subtyping them at the highest possible level of resolution. Because wgMLST harnesses the full power of Illumina sequencing for characterizing isolates, it can be used for discriminating between closely-related isolates that have diversified over a short timeframe. This is highly relevant during outbreak investigations, for tracking the origin of contamination, and precise assessment of divergence dates. This discriminatory power is not achieved with cgMLST because it only assesses differences in the core genome and has been shown to provide fewer allelic differences in comparison to wgMLST (10). Further, cgMLST schemes are mostly composed of slowly evolving genes. Previous studies on *Lm* genomes have estimated the evolution rate of cgMLST types to be around 0.2 alleles per year, indicating that cgMLST-based typing is insufficient for discriminating isolates which have diverged over short timeframes (31). However, use of a well-defined set of species-wide conserved genes makes cgMLST more stable and suitable for robust comparisons of distantly related isolates. Typically, cgMLST is sufficient for routine epidemiological surveillance such as identification of clonal groups and discrimination of outbreak strains from epidemiologically unrelated strains. Haplo-ST can perform both core-genome and whole-genome MLST because its database incorporates genes in the core-genome (the *Lm* cgMLST scheme developed by Institut Pasteur) together with accessory genes in the pan-genome of *Lm*. Additionally, it can be used for inferring biological properties such as virulence, antibiotic-resistance, stress tolerance and phenotypic predictions like serotypes by profiling genes linked to these properties. The wgMLST scheme in Haplo-ST can also be expanded to include genotypic variation in future *Lm* isolates by updating the locally installed BIGSdb-*Lm* database housed within this platform. This can include multi-copy and accessory genes which may arise through recombination and whose detection may become important for pathogen surveillance.

Unlike SNP-based genotyping which uses individual SNPs as units of comparison, cg/wgMLST counts different types of variants within one coding region as a single allelic change. This concept covers the conflicting signals of horizontal and vertical transfer of genetic material as a single evolutionary event and classifies WGS data as a set of allele identifiers, thereby enabling easy storage of a stable nomenclature within a database and making comparisons of wgMLST profiles faster. Nonetheless, this also leads to a loss of resolution as it obscures the extent of dissimilarity between non-identical alleles. Thus, the technical performance of wgMLST along with its amenability to standardization is accompanied by a loss in specificity, as minimum spanning trees constructed using sequence types are fully connected, failing to effectively split isolate populations into clonal complexes (36). This becomes problematic as allele-based subtyping alone does not provide sufficient information for delineating outbreaks; it is therefore critical to complement it with whole-genome based phylogenetic clustering for accessing relationships between isolates (37). Recent studies have shown that although wgMLST-based dendrograms are comparable to SNP-based phylogenies in identifying clades of closely related isolates with a recent common ancestor, they differ from each other with respect to the placement of isolates within clonal groups, where branches in SNP-based phylogenies are not supported by greater than 90% bootstrap support (10). This emphasizes the importance of constructing phylogenies with confidence measures such as bootstrap support, which is unfortunately not feasible with wgMLST-based dendrograms. Haplo-ST has the advantage of not only providing wgMLST profiles, but also provides corresponding allele sequences assembled for each isolate. While allelic profiles can be used for constructing dendrograms from allelic similarity type matrices, allele sequences can be concatenated and used for constructing cg/wgMLST-based phylogenies using a variety of models of molecular evolution and obtaining bootstrap support values. Moreover, our tool can detect paralogous genes which when ignored, can lead to the construction of biased phylogenies. Thus, analysis provided by Haplo-ST, when combined with detailed epidemiological evidence, isolate metadata and appropriate interpretation allows for routine surveillance of *Lm*, accurate source-tracking of contaminating strains, elucidation of transmission pathways and ultimately helps in devising better intervention strategies in food safety monitoring programs.

Our approach was evaluated for its usability in characterizing and determining relatedness within two groups of *Lm* isolates; one group representing isolates present in the natural environment and the other from poultry further processing facilities. This enabled us to decipher the phylogenetic relatedness of *Lm* isolates which shows clear delineation between lineages in both isolate groups. A majority of isolates in the natural environment and food facilities belonged to lineage II, which is consistent with previous studies (21). Further, the lineage of 11% isolates could not be identified with lineage-specific probes. All of these were identified using our methods, including 2% that belonged to other species. Moreover, we did not find significant differences in the distribution of isolates belonging to different lineages in terms of their phenotypes (persistent/transient) and origin (sampling sites). However, it is curious that we found no lineage III isolates in the processing plant samples but 17% in the natural environment.

*Lm* is a foodborne pathogen that is prevalent in the natural environment. Its ability to colonize and persist in food processing environments increases the risk of contaminating ready-to-eat (RTE) food, often leading to outbreaks of listeriosis. Hence, understanding the genetic determinants associated with its adaptation and persistence in food processing environments is of paramount importance for developing targeted intervention strategies, and the typing of *Lm* plays a crucial role in such investigations. Another advantage of the wgMLST approach is that it lends itself to analyses that help to form hypotheses on the mechanisms of segregation of isolates.

We used Haplo-ST to type and identify loci with significant genetic variation between isolates obtained from the natural environment and poultry processing facilities. Our analysis revealed 111 significantly differentiated loci which may be involved in helping *Lm* to adapt to high stress conditions within food processing environments, thereby increasing its risk of contaminating food. Unlike transient strains, which are frequently introduced into food facilities from the natural environment and easily removed with regular sanitation shifts, persistent strains have been reported to have enhanced capacity to adapt and survive in food production chains and are difficult to eradicate. Thus, we also used our tool to characterize and detect loci with high genomic differentiation between transient and persistent strains. We obtained 102 highly differentiated loci potentially enriched for the ‘persistent’ phenotype. Of these, 21 loci were common with the 111 loci we previously identified as potentially contributing towards adaptation in food processing facilities (Table 2). These loci were considered to be significantly enriched for adaptation and were related to metabolism (*lmo0875, lmo2650, lmo1336, lmo1817, lmo1464, lmo2640*), transport (*lmo0875, lmo2650, lmo1210, lmo2383, lmo1960, lmo1205*), tRNA and ribosome biogenesis (*lmo1949, lmo2078, lmo1294*), biosynthesis of secondary metabolites (*lmo1294, lmo2640*), translation (*lmo2548, lmo2073*), and oxidative stress (*lmo0964*). We also found that out of the 102 loci differentiated for persistence, three genes (*lmo1699, lmo0692, lmo2020*) were found to be associated with chemotaxis, a process that plays a role in niche localization (38). Several studies have shown the presence of a five gene stress survival islet, *SSI-1* to contribute to the growth of *Lm* under suboptimal conditions like low pH and high salt concentrations (39, 40). Our analyses found *SSI-1* in a higher fraction of isolates (93%) from processing plants compared to the natural environment (17%). Other studies report resistance to quaternary ammonium compounds like benzalkonium chloride (BC) in persistent strains (41). BC is commonly used as an agri-food sanitizer and resistance to it is provided by the gene cassette *bcrABC*, in which *bcrAB* codes for the small multidrug resistance protein family transportera and *bcrC* codes for a transcriptional factor. Our subtyping results are in agreement with this; *bcrABC* was present in 72% of the isolates obtained from the effluents, but absent in isolates obtained from the natural environment. Among isolates collected from effluents, *bcrABC* was associated with a higher proportion of persistent strains (54%) when compared to transient strains (18%).

Our approach does, however, have a few limitations. Although the locally installed database within our platform is expandable to accommodate future genetic diversity in *Lm*, it requires frequent manual upgrades as new alleles and genes become available. With the recent accessibility of BIGSdb-*Lm* at Pasteur Institut through RESTful API, this drawback can be resolved by making minor modifications to our pipeline which will allow the tool to interrogate the server at Pasteur Institut directly instead of calling alleles locally. Secondly, our approach is gene-centric and characterizes differences only in protein-coding genes; therefore, genetic variation in other genomic regions like pseudogenes and intergenic regions are not accounted for. Additionally, the use of short reads may produce faulty assemblies of accessory genes and repeat regions. With the decreasing costs and increased popularity of third-generation sequencing instruments, these limitations can be overcome with development of appropriate sequence assembly algorithms. Thus, the power of fully assembled genomes remain yet to be exploited. Nevertheless, the current wgMLST approach will be stable over time as new genes are added and maintain backwards compatibility with classical seven-gene MLST schemes.

The greatest advantage of Haplo-ST is that this platform is flexible and not limited to profiling of *Listeria monocytogenes* alone. It can be adapted to provide molecular characterization for any haploid organism, with the installation of an organism-specific gene database with associated allelic nomenclature, along with minor changes to the script that automates the pipeline. Furthermore, users are not limited to using publicly available gene databases because BIGSdb can accommodate any custom user-provided database.

## MATERIALS AND METHODS

### Development of Haplo-ST for wgMLST profiling of *Lm* strains

We developed Haplo-ST to analyze wgMLST for *Lm* (Fig. 1). This tool takes in raw WGS reads for each *Lm* isolate and uses the FASTX-Toolkit v0.0.14 (42) to clean them according to user-specified parameters. It then uses YASRA v2.33, (available at https://github.com/aakrosh/YASRA; 43) to assemble genes across loci by mapping to reference genes. We selected YASRA for assembling genes because YASRA is a comparative assembler which uses a template to guide the assembly of a closely related target sequence, and can accommodate high rates of polymorphism between the template and target (43). Hence, this assembler can be used to assemble an allelic variant of a gene by mapping to a reference sequence, even when the target allele has diverged considerably from the reference gene sequence. Next, a local installation of the BIGSdb-*Lm* database (available at http://bigsdb.pasteur.fr/listeria, 44) is used by Haplo-ST to assign allelic profiles to the genes assembled with YASRA, thus generating a wgMLST profile for each isolate. The BIGSdb-*Lm* database contains allelic profiles of 2554 *Lm* genes obtained from BIGSdb-*Lm* as of 2^nd^ June 2017. This pipeline has been automated with a Perl script and made portable by installation of all software dependencies along with a local installation of the BIGSdb-*Lm* database within a Linux Virtual Machine (VM). Haplo-ST has been made available at https://github.com/swarnalilouha/Haplo-ST. In addition to generating wgMLST profiles, Haplo-ST also outputs the list of gene sequences assembled for each isolate. Because BIGSdb-*Lm* can identify all paralogs associated with a query gene sequence as ‘exact matches’, our tool has also been automated to output a list of paralogs identified for each isolate.

### Sensitivity of Haplo-ST

ART v2.5.8 (45) was used to simulate WGS reads for two reference genomes of *Lm*, EGD-e (NCBI accession number NC_003210.1) and Strain 4b F2365 (NCBI accession number NC_002973.6). The simulated WGS reads were of two different lengths (150 bp and 250 bp); and different qualities, one set of reads with high quality throughout the read length and the other with degrading quality over the length of the read. In total, 8 sets of simulated WGS reads were processed through Haplo-ST to generate 8 wgMLST profiles. Four of these wgMLST profiles were obtained from simulated reads generated from the *Lm* EGD-e reference genome. Each of these 4 profiles were compared to the allelic profiles of annotated genes in EGD-e. The other four wgMLST profiles were obtained from reads derived from the Strain 4b F2365 reference genome. These were compared to the allelic profiles of annotated genes in F2365. For each comparison, we calculated the percentage of genes correctly typed by Haplo-ST. Finally, we calculated the average sensitivity over eight comparisons.

### Dependency of Haplo-ST on sequencing depth

To determine the levels of genome sequence coverage necessary for efficient whole-genome sequence typing, synthetic reads were simulated from the *Lm* EGD-e reference genome with ART v2.5.8 for different sequencing depths ranging from 5× - 120× and typed with Haplo-ST (performed in triplicate). For each sequencing depth, the allelic profiles typed by our tool were compared to allelic profiles of annotated genes from the *Lm* EGD-e reference genome. Finally, for each comparison, we calculated: (i) the number of genes correctly typed, (ii) the number of genes assigned an erroneous allele ID, (iii) the number of genes partially assembled and (iv) the number of genes missing an allele ID assignment by Haplo-ST.

### Analysis of *Lm* strains collected from the natural environment and poultry processing plants

#### Lm Isolate collection, DNA extraction and sequencing

*Lm* isolates obtained from the natural environment were cultured from water and sediment samples collected at 16 locations in the South Fork Broad River watershed located in Northeast Georgia (46). Sampling locations were selected based on predominant land use by the National Land Cover Database and on-the-ground surveys. Samples were collected from 6 sites designated as agricultural/pastoral; 7 sites as forested; 2 sites as impacted by water pollution control plants (WPCP) and 1 site classified as mixed-use. *Lm* isolates obtained from poultry processing plants were sampled from different locations within the poultry processing plants at different time periods (12, 47). Some of these isolates were repeatedly isolated from multiple sites in the plants over an extended period of time and were designated as ‘persistent’ types (based on actA-sequence subtyping); other isolates sporadically isolated from the food processing facilities were classified as ‘transient’ strains. Each colony of *Lm* isolate cultured from the samples was inoculated into 5ml of tryptic soy broth and grown overnight at 35 °C. DNA was extracted using the UltraClean^®^ Microbial DNeasy Kit (Qiagen, Venlo, The Netherlands) according to manufacturer’s instructions. Sequencing libraries were prepared using the Nextera XT DNA Library Preparation Kit (Illumina, San Diego, USA). Genomic DNA of each isolate was sequenced using the Illumina MiSeq platform to obtain paired-end 150 or 250 bp reads. This effort yielded WGS data for a total of 171 *Lm* isolates obtained from the natural environment (NCBI BioProject Accession: PRJNA605751) and 162 *Lm* isolates obtained from poultry processing plants (NCBI BioProject Accession: PRJNA606479). These were then processed using Haplo-ST.

#### wgMLST profiling of Lm isolates with Haplo-ST

WGS data for *Lm* isolates was first checked for quality with FastQC v0.11.4 (48). The raw data was then cleaned with the FASTX-Toolkit v0.0.14 incorporated within Haplo-ST. User-specified parameters were used to perform three successive cleaning steps with FASTA/Q Trimmer, FASTQ Quality Trimmer and FASTQ Quality Filter tools of the FASTX-Toolkit. Reads were trimmed to remove all bases with a Phred quality score of < 20 from both ends, and filtered such that 90% of bases in the clean reads had a quality of at least 20. After trimming and filtering, all remaining reads with lengths of < 50 bp were filtered out. Next, the cleaned reads were assembled into gene sequences by mapping to reference genes with YASRA. While assembling genes across loci, all assemblies having a length of less than 89% the length of the corresponding reference gene were removed. This is because our examination of the lengths of all 2554 genes and their respective alleles in the BIGSdb-*Lm* database revealed that alleles of a gene can have different lengths, which ranges from 0.89 - 1.09 times the length of the reference gene. This ‘length criteria’ for filtering assembled genes has been provided as a user-specified parameter in the Perl script that automates Haplo-ST. The value for this parameter can be adjusted if the BIGSdb-*Lm* database is updated to include more genes or alleles, or if only a subset of genes is used for allelic profiling. Finally, assembled genes were assigned allele ID’s with BIGSdb-*Lm* and wgMLST profiles were generated for each isolate.

#### Identification of paralogous genes

We identified paralogous genes in our dataset using two approaches. In the first approach, Haplo-ST uses BIGSdb-*Lm*’s ability to identify paralogs and outputs a list of paralogs for each isolate. To verify that all paralogs were correctly identified with BIGSdb-*Lm*, we used a second approach to detect paralogs present within the BIGSdb-*Lm* database. First a local BLAST database was created with all 2554 genes and their corresponding alleles present in the BIGSdb-*Lm* database using BLAST+ v2.2.29. Next, BLAST searches of all genes and their respective alleles were made against the local BLAST database (File S1). Custom Perl scripts were used to identify genes having an exact sequence match to another gene in the database and all such matches were listed as paralogs.

#### Construction of phylogenetic trees and evaluation of lineage-specific association

The list of genes assembled for each isolate with Haplo-ST were filtered to remove paralogous genes. The final filtered assemblies for each group of isolates (the first group obtained from the natural environment and the second group obtained from poultry processing plants) were used to create concatenated multiple sequence alignments (MSA) with Phyluce v1.5.0 (49). Several scripts were used to create MSA’s for each isolate group. First, a custom Perl script was used to convert the assembled gene sequences into a format suitable for use with Phyluce. Second, the ‘phyluce_align_seqcap_align’ script was used to align genes across loci for all isolates within a group and the alignment was trimmed for ragged edges. The summary statistics of alignments for both isolate groups were checked with the script ‘phyluce_align_get_align_summary_data’ and cleaned for locus names with ‘phyluce_align_remove_locus_name_from_nexus_lines’. The dataset for each isolate group was then culled to reach a 95% level of completeness with ‘phyluce_align_get_only_loci_with_min_taxa’. The 95% complete data matrix was converted into phylip files with ‘phyluce_align_format_nexus_files_for_raxml’ and phylogenetic trees were constructed with FastME v2.1.5 (50). The substitution model used by FastME was ‘p-distance’ and the BioNJ algorithm was used to compute a tree from the distance matrix (File S2). A total of 500 bootstrap replicates were computed to provide support to the internal branches of each of the phylogenies.

*Lm* isolates were classified into lineages (I to IV) based on a targeted multilocus genotyping approach (TMLGT) in which six genomic regions were coamplified in a multiplexed PCR and used as templates for allele-specific primer extension using lineage-specific probes (51). Lineage-specific correlation between groups of isolates was tested with Fisher’s exact test at *P* = 0.05.

Phylogenetic trees were visualized and annotated with iTOL v3 (52). For better visualization, all phylogenetic trees were converted to circular format and lineage classification for isolates was displayed by coloring internal branches. The annotations for the source and type of isolates were displayed in outer external rings.

#### Analysis of genetic variation

To obtain measures of genetic differentiation, we used the wgMLST profiles from Haplo-ST and performed Analysis of Molecular Variance (AMOVA) in Arlequin v3.5.2 (53). First, paralogous loci were removed from the raw wgMLST profiles. Next, new alleles not defined in the BIGSdb-*Lm* database and reported as ‘closest matches’ to existing alleles in the wgMLST profiles were assigned custom allele ID’s with in-house python scripts. Finally, AMOVA was separately performed at two levels: (i) among groups of isolates obtained from the natural environment and poultry processing plants, and (ii) among groups of transient and persistent strains obtained from the poultry processing plants. For each level of analysis, loci with < 10% missing data in the wgMLST profiles were used. Fifty thousand permutations were used to determine significance of variance components. In addition to AMOVA which calculates the global F_ST_ for all loci within a group of isolates, we also performed a locus by locus AMOVA which computes F_ST_ indices for each locus separately, for both levels of analysis. The upper 5% of the distribution of F_ST_ values was chosen as the threshold for loci with significant genetic diversity.

## ACKNOWLEDGEMENTS

This research was supported by funding from USDA Agricultural Research Service Project Number 6040-32000-009-00-D. We thank USDA and FSIS for providing us with *Listeria monocytogenes* whole-genome sequencing samples for our work. We also thank Yecheng Huang for assistance with a local installation of the BIGSdb database. The high-performance computing cluster at Georgia Advanced Computing Resource Center (GACRC) at the University of Georgia provided computational infrastructure and technical support throughout the work.

## SUPPLEMENTAL MATERIAL

**File S1:** Script for making BLAST searches of all genes and their respective alleles against a local BLAST database.

**File S2:** Script for constructing phylogenetic trees with FastME.

**File S3:** List of 133 paralogous genes identified in our dataset.

**File S4:** Inaccurate characterization by BIGSdb-*Lm* when it recognizes non-paralogous genes as ‘exact matches’.

**File S5:** List of 111 loci with the highest F_ST_ values (top 5% of F_ST_ distribution) having considerable genetic variation between isolates obtained from the natural environment and poultry processing plants.

**File S6:** List of 102 loci (upper 5% of F_ST_ distribution) having the most genetic divergence between the transient and persistent strains.

**File S7:** Whole-genome MLST profiles of *Lm* isolates generated by Haplo-ST.

**File S8:** Whole-genome MLST profiles of *Lm* isolates with new alleles assigned custom allele-IDs.

## REFERENCES

1. Bennion JR, Sorvillo F, Wise ME, Krishna S, Mascola L. 2008. Decreasing listeriosis mortality in the United States, 1990-2005. Clin Infect Dis 47:867–74.

2. Scallan E, Hoekstra RM, Angulo FJ, Tauxe RV, Widdowson MA, Roy SL, Jones JL, Griffin PM. 2011. Foodborne illness acquired in the United States--major pathogens. Emerg Infect Dis 17:7–15.

3. USDA ERS. 2014. Cost estimates of foodborne illnesses. Economic Re- search Service, US Department of Agriculture, Washington, DC. Available at: https://www.ers.usda.gov/data-products/cost-estimates-of-foodborne-illnesses.aspx.

4. Den Bakker HC, Didelot X, Fortes ED, Nightingale KK, Wiedmann M. 2008. Lineage specific recombination rates and microevolution in *Listeria monocytogenes*. BMC Evol Biol 8:277.

5. Haase JK, Didelot X, Lecuit M, Korkeala H, L. monocytogenes MLST Study Group; Achtman M. 2014. The ubiquitous nature of *Listeria monocytogenes* clones: a large-scale Multilocus Sequence Typing study. Environ Microbiol 16:405–16.

6. Painset A, Björkman JT, Kiil K, Guillier L, Mariet JF, Félix B, Amar C, Rotariu O, Roussel S, Perez-Reche F, Brisse S, Moura A, Lecuit M, Forbes K, Strachan N, Grant K, Møller-Nielsen E, Dallman TJ. 2019. LiSEQ - whole-genome sequencing of a cross-sectional survey of *Listeria monocytogenes* in ready-to-eat foods and human clinical cases in Europe. Microb Genom 5: e000257.

7. Hyden P, Pietzka A, Lennkh A, Murer A, Springer B, Blaschitz M, Indra A, Huhulescu S, Allerberger F, Ruppitsch W, Sensen CW. 2016. Whole genome sequence-based serogrouping of *Listeria monocytogenes* isolates. J Biotechnol 235:181–6.

8. Orsi RH, Borowsky ML, Lauer P, Young SK, Nusbaum C, Galagan JE, Birren BW, Ivy RA, Sun Q, Graves LM, Swaminathan B, Wiedmann M. 2008. Short-term genome evolution of *Listeria monocytogenes* in a non-controlled environment. BMC Genom 9:539.

9. Carpentier B, Cerf O. 2011. Review–persistence of *Listeria monocytogenes* in food industry equipment and premises. Int J Food Microbiol 145:1–8.

10. Jagadeesan B, Baert L, Wiedmann M, Orsi RH. 2019. Comparative Analysis of Tools and Approaches for Source Tracking *Listeria monocytogenes* in a Food Facility Using Whole-Genome Sequence Data. Front Microbiol 10:947.

11. Kuhn M, Goebel W. 2007. Molecular virulence determinants of *Listeria monocytogenes*, p 111–155. In Ryser ET, Marth EH (ed), Listeria, listeriosis and food safety, 3rd ed, CRC Press Taylor and Francis Group, Boca Raton, FL.

12. Berrang ME, Meinersmann RJ, Frank JF, Smith DP, Genzlinger LL. 2005. Distribution of *Listeria monocytogenes* subtypes within a poultry further processing plant. J Food Prot 68:980–985.

13. Moorman M, Pruett P, Weidman M. 2010. Value and Methods for Molecular Subtyping of Bacteria, p 157–175. In Kornacki JL (ed), Principles of Microbiological Troubleshooting in the Industrial Food Processing Environment, 1st ed, Springer Science Business Media, New York, NY.

14. Swaminathan B, Barrett T, Hunter SB, Tauxe RV, CDC PulseNet Task Force. 2001. PulseNet: The molecular subtyping network for foodborne bacterial disease surveillance, United States. Emerg Infect Dis 7:382–389.

15. Ruppitsch W, Pietzka A, Prior K, Bletz S, Fernandez HL, Allerberger F, Harmsen D, Mellmann A. 2015. Defining and evaluating a core genome MLST scheme for whole genome sequence-based typing of *Listeria monocytogenes*. J Clin Microbiol 53:2869–76.

16. Henri C, Félix B, Guillier L, Leekitcharoenphon P, Michelon D, Mariet JF, Aarestrup FM, Mistou MY, Hendriksen RS, Roussel S. 2016. Population Genetic Structure of *Listeria monocytogenes* Strains as Determined by Pulsed-Field GelElectrophoresis and Multilocus Sequence Typing. Appl Environ Microbiol. 82:5720–8.

17. Ragon M, Wirth T, Hollandt F, Lavenir R, Lecuit M, Monnier Le A, Brisse S. A new perspective on *Listeria Monocytogenes* evolution. 2008. PLoS Pathog 4:e1000146.

18. Orsi RH, den Bakker HC, Wiedmann M. 2011. *Listeria monocytogenes* lineages: genomics, evolution, ecology, and phenotypic characteristics. Int J Med Microbiol 301:79–96.

19. Meinersmann RJ, Phillips RW, Wiedmann M, Berrang ME. 2004. Multilocus Sequence Typing of *Listeria Monocytogenes* by Use of Hypervariable Genes Reveals Clonal and Recombination Histories of Three Lineages. Appl Environ Microbiol 70:2193–203.

20. Pirone-Davies C, Chen, Y, Pightling A, Ryan G, Wang Y, Yao K, Hoffmann M, Allard MW. 2018. Genes significantly associated with lineage II food isolates of *Listeria monocytogenes*. BMC Genomics 19:708.

21. Dreyer M, Aguilar-Bultet L, Rupp S, Guldimann C, Stephan R, Schock A, Otter A, Schüpbach G, Brisse S, Lecuit M, Frey J, Oevermann A. 2016. *Listeria monocytogenes* sequence type 1 is predominant in ruminant rhombencephalitis. Sci Rep 6:36419.

22. Jackson BR, Tarr C, Strain E, Jackson KA, Conrad A, Carleton H, Katz LS, Stroika S, Gould LH, Mody RK, Silk BJ, Beal J, Chen Y, Timme R, Doyle M, Fields A, Wise M, Tillman G, Defibaugh-Chavez S, Kucerova Z, Sabol A, Roache K, Trees E, Simmons M, Wasilenko J, Kubota K, Pouseele H, Klimke W, Besser J, Brown E, Allard M, Gerner-Smidt P. 2016. Implementation of Nationwide Real-time Whole-genome Sequencing to Enhance Listeriosis Outbreak Detection and Investigation. Clin Infect Dis 63:380–386.

23. Moura A, Tourdjman M, Leclercq A, Hamelin E, Laurent E, Fredriksen N, Van Cauteren D, Bracq-Dieye H, Thouvenot P, Vales G, Tessaud-Rita N, Maury MM, Alexandru A, Criscuolo A, Quevillon E, Donguy MP, Enouf V, de Valk H, Brisse S, Lecuit M. 2017. Real-Time Whole-Genome Sequencing for Surveillance of *Listeria monocytogenes*, France. Emerg Infect Dis 23:1462–1470.

24. Halbedel S, Prager R, Fuchs S, Trost E, Werner G, Flieger A. Whole-Genome Sequencing of Recent *Listeria monocytogenes* Isolates from Germany Reveals Population Structure and Disease Clusters. 2018. J Clin Microbiol 56:e00119–18.

25. Kvistholm Jensen A, Nielsen EM, Björkman JT, Jensen T, Müller L, Persson S, Bjerager G, Perge A, Krause TG, Kiil K, Sørensen G, Andersen JK, Mølbak K, Ethelberg S. 2016. Whole-genome Sequencing Used to Investigate a Nationwide Outbreak of Listeriosis Caused by Ready-to-eat Delicatessen Meat, Denmark, 2014. Clin Infect Dis 63:64–70.

26. Kwong JC, Mercoulia K, Tomita T, Easton M, Li HY, Bulach DM, Stinear TP, Seemann T, Howden BP. 2016. Prospective Whole-Genome Sequencing Enhances National Surveillance of *Listeria monocytogenes*. J Clin Microbiol 54:333–342.

27. Katz LS, Griswold T, Williams-Newkirk AJ, Wagner D, Petkau A, Sieffert C, Domselaar GV, Deng X, Carleton HA. 2017. A comparative analysis of the Lyve-SET phylogenomics pipeline for genomic epidemiology of foodborne pathogens. Front Microbiol 8:375.

28. Henri C, Leekitcharoenphon P, Carleton HA, Radomski N, Kaas RS, Mariet JF, Felten A, Aarestrup FM, Gerner Smidt P, Roussel S, Guillier L, Mistou MY, Hendriksen RS. 2017. An Assessment of Different Genomic Approaches for Inferring Phylogeny of *Listeria monocytogenes*. Front Microbiol 8:2351.

29. Pightling AW, Petronella N, Pagotto F. 2015. The *Listeria monocytogenes* core -genome sequence typer (LmCGST): a bioinformatics pipeline for molecular characterization with next generation sequence data. BMC Microbiol 15:224.

30. Chen Y, Gonzalez-Escalona N, Hammack TS, Allard MW, Strain EA, Brown EW. 2016. Core Genome Multilocus Sequence Typing for Identification of Globally Distributed Clonal Groups and Differentiation of Outbreak Strains of *Listeria monocytogenes*. Appl Environ Microbiol 82:6258–6272.

31. Moura A, Criscuolo A, Pouseele H, Maury MM, Leclercq A, Tarr C, Björkman JT, Dallman T, Reimer A, Enouf V, Larsonneur E, Carleton H, Bracq-Dieye H, Katz LS, Jones L, Touchon M, Tourdjman M, Walker M, Stroika S, Cantinelli T, Chenal-Francisque V, Kucerova Z, Rocha EPC, Nadon C, Grant K, Nielsen EM, Pot B, Gerner-Smidt P, Lecuit M, Brisse S. 2016. Whole genome-based population biology and epidemiological surveillance of *Listeria monocytogenes*. Nat Microbiol 2:16185.

32. Pietzka A, Allerberger F, Murer A, Lennkh A, Stöger A, Cabal Rosel A, Huhulescu S, Maritschnik S, Springer B, Lepuschitz S, Ruppitsch W, Schmid D. 2019. Whole Genome Sequencing Based Surveillance of *L. monocytogenes* for Early Detection and Investigations of Listeriosis Outbreaks. Front Public Health 7:139.

33. Wang J, Ray AJ, Hammons SR, Oliver HF. 2015. Persistent and transient *Listeria monocytogenes* strains from retail deli environments vary in their ability to adhere and form biofilms and rarely have inlA premature stop codons. Foodborne Pathog Dis 12:151–8.

34. Autio T, Keto-Timonen R, Lundén J, Björkroth J, Korkeala H. 2003. Characterization of persistent and sporadic *Listeria monocytogenes* strains by pulsed-field electrophoresis (PFGE) and amplified fragment length polymorphism (ALFP). Syst Appl Microbiol 26:539–45.

35. Maiden MC, Jansen van Rensburg MJ, Bray JE, Earle SG, Ford SA, Jolley KA, McCarthy ND. 2013. MLST revisited: the gene-by-gene approach to bacterial genomics. Nat Rev Microbiol 11:728–36.

36. Feil EJ, Li BC, Aanensen DM, Hanage WP, Spratt BG. 2004. eBURST: inferring patterns of evolutionary descent among clusters of related bacterial genotypes from multilocus sequence typing data. J Bacteriol 186:1518–30.

37. Chen Y, Luo Y, Carleton H, Timme R, Melka D, Muruvanda T, Wang C, Kastanis G, Katz LS, Turner L, Fritzinger A, Moore T, Stones R, Blankenship J, Salter M, Parish M, Hammack TS, Evans PS, Tarr CL, Allard MW, Strain EA, Brown EW. 2017. Whole Genome and Core Genome Multilocus Sequence Typing and Single Nucleotide Polymorphism Analyses of *Listeria monocytogenes* Isolates Associated with an Outbreak Linked to Cheese, United States, 2013. Appl Environ Microbiol 83:e00633–17.

38. Casey A, Fox EM, Schmitz-Esser S, Coffey A, McAuliffe O, Jordan K. 2014. Transcriptome analysis of *Listeria monocytogenes* exposed to biocide stress reveals a multi-system response involving cell wall synthesis, sugar uptake, and motility. Front Microbiol 5:68.

39. Ryan S, Begley M, Hill C, Gahan CGM. 2010. A Five-Gene Stress Survival Islet (SSI-1) That Contributes to the Growth of *Listeria Monocytogenes* in Suboptimal Conditions. J Appl Microbiol 109:984–95.

40. Gómez D, Azón E, Marco N, Carramiñana JJ, Rota C, Ariño A, Yangüela J. 2014. Antimicrobial Resistance of *Listeria Monocytogenes* and *Listeria Innocua* from Meat Products and Meat-Processing Environment. Food Microbiol 42:61–5.

41. Cherifi T, Carrillo C, Lambert D, Miniaï I, Quessy S, Larivière-Gauthier G, Blais B, Fravalo P. 2018. Genomic Characterization of *Listeria Monocytogenes* Isolates Reveals That Their Persistence in a Pig Slaughterhouse Is Linked to the Presence of Benzalkonium Chloride Resistance Genes. BMC Microbiol 18:220.

42. Hannon GJ. 2010. FASTX-Toolkit, FASTQ/A short-reads pre-processing tools. Repository http://hannonlab.cshl.edu/fastx_toolkit

43. Ratan A. 2009. Assembly algorithms for next generation sequence data. Ph.D. dissertation, The Pennsylvania State University.

44. Jolley KA, and Maiden MC. 2010. BIGSdb:scalable analysis of bacterial genome variation at the population level. BMC Bioinform 11:595.

45. Huang W, Li L, Myers JR, Marth GT. 2012. ART: A Next-Generation Sequencing Read Simulator. Bioinformatics 28:593–4.

46. Bradshaw JK, Snyder BJ, Oladeinde A, Spidle D, Berrang ME, Meinersmann RJ, Oakley B, Sidle RC, Sullivan K, Molina M. 2016. Characterizing relationships among fecal indicator bacteria, microbial source tracking markers, and associated waterborne pathogen occurrence in stream water and sediments in a mixed land use watershed. Water Res 101:498–509.

47. Berrang ME, Meinermann RJ, Frank JF, Ladely SR. 2010. Colonization of a newly constructed commercial chicken further processing plant with *Listeria monocytogenes*. J Food Prot 73:286–291.

48. Andrews S. 2010. FastQC: a quality control tool for high throughput sequence data. Repository http://www.bioinformatics.babraham.ac.uk/projects/fastqc

49. Faircloth BC. 2016. PHYLUCE is a software package for the analysis of conserved genomic loci. Bioinformatics 32:786–8.

50. Lefort V, Desper R, Gascuel O. 2015. FastME 2.0: A Comprehensive, Accurate, and Fast Distance-Based Phylogeny Inference Program. Mol Biol Evol 32:2798–800.

51. Ward TJ, Usgaard T, Evans P. 2010. A targeted multilocus genotyping assay for lineage, serogroup, and epidemic clone typing of *Listeria monocytogenes*. Appl Environ Microbiol 76:6680–4.

52. Letunic I, Bork P. 2016. Interactive tree of life (iTOL) v3: an online tool for the display and annotation of phylogenetic and other trees. Nucleic Acids Res 44:W242–5.

53. Excoffier L, Lischer HEL. 2010. Arlequin suite ver 3.5: A new series of programs to perform population genetics analyses under Linux and Windows. Mol Ecol Resour 10:564–567.

